# Neural mechanisms of flexible perceptual inference

**DOI:** 10.1101/2025.02.05.636696

**Authors:** John Schwarcz, Jan Bauer, Haneen Rajabi, Gabrielle Marmur, Eran Lottem, Jonathan Kadmon

## Abstract

What seems obvious in one context can take on an entirely different meaning if that context shifts. While context-dependent inference has been widely studied, a fundamental question remains: how does the brain simultaneously infer both the meaning of sensory input and the underlying context itself, especially when the context is changing? Here, we study flexible perceptual inference—the ability to adapt rapidly to implicit contextual shifts without trial and error. We introduce a novel change-detection task in dynamic environments that requires tracking latent state and context. We find that mice exhibit first-trial behavioral adaptation to latent context shifts driven by inference rather than reward feedback. By deriving the Bayes-optimal policy under a partially observable Markov decision process, we show that rapid adaptation emerges from sequential updates of an internal belief state. In addition, we show that artificial neural networks trained via reinforcement learning achieve near-optimal performance, implementing Bayesian inference-like mechanisms within their recurrent dynamics. These networks develop flexible internal representations that enable real-time adjustments to the inference model. Our findings establish flexible perceptual inference as a core principle of cognitive flexibility, offering computational and neural-mechanistic insights into adaptive behavior in uncertain environments.

## 1 Introduction

As dusk settles over a field, a mouse hesitates at the edge of its burrow, torn between hunger and caution. Every rustle, shadow, and scent carries ambiguity—harmless background noise or a lurking predator? A flicker in the sky—owl or drifting cloud? Each cue is uncertain, yet survival hinges on interpreting them swiftly and accurately. Crucially, the mouse’s decisions are shaped not only by immediate evidence, but also by an inferred context: Is the wind in the underbrush stronger than usual, making rustling sounds less reliable? Are the shadows deeper tonight, obscuring movement? Rather than passively accumulating sensory evidence, the mouse must continuously update its internal model of the world, refining both its beliefs about what is happening and how incoming information should be weighted and interpreted.

This dual challenge—extracting meaning from uncertain inputs while simultaneously inferring the latent structure that shapes their interpretation—is fundamental to perception and decision-making. The brain must solve it efficiently and continuously. This process, which we term **flexible perceptual inference**, enables organisms to rapidly adapt to changing conditions, resolve uncertainty, and make informed decisions without relying solely on trial-and-error learning. This cognitive flexibility is as crucial for a mouse navigating the wilderness as it is for any intelligent system operating in a complex and unpredictable world^1^.

The challenge of probabilistic reasoning in uncertain environments is best captured by Bayesian decision theory, which provides a principled framework for perceptual inference. This framework formalizes how optimal agents update their beliefs in response to uncertain observations^2–4^. Empirical studies have shown that human and animal behavior is often aligned with Bayesian inference, particularly in perceptual decision-making tasks where organisms integrate sequential sensory evidence to make decisions under uncertainty^5–8^.

A common framework for modeling the accumulation of perceptual evidence is the drift-diffusion model (DDM)^9–12^, which approximates Bayesian inference under different conditions^13,14^. However, traditional DDM formulations assume a fixed and known context, leaving open the question of how real-time inference adapts to latent contextual uncertainty. Much of the existing work on learning in changing environments has focused on how organisms detect and adjust to changes in volatility^15–17^, often by modifying learning rates or belief update rules in hierarchical Bayesian models^18–20^.

Existing studies on contextual evidence integration typically fall into three main categories. The first examines how neural circuits and artificial networks perform context-dependent integration but assume an explicit externally provided context signal^21–24^. The second asks how humans and animals adapt decision making in dynamic environment by discounting old evidence^25–28^. The third category considers contextual inference, where the context must be inferred from observations^29–31^. However, the mechanisms that allow the brain to infer an implicit context and adapt the dynamics of evidence accumulation simultaneously remain poorly understood^32^.

Crucially, distinguishing between these mechanisms is challenging, as behavior alone does not uniquely constrain the underlying neural computations^33–37^. Multiple evidence accumulation models can generate behavior that appears consistent with Bayesian inference, making it difficult to infer how the brain implements flexible perceptual inference. Resolving this ambiguity requires moving beyond behavioral signatures to examine how neural circuits represent, update, and adapt decision variables in response to changing contexts^38–41^.

To address these open questions, we investigate how neural circuits perform *flexible perceptual inference*—how they simultaneously infer both latent states and context from ambiguous sensory input and dynamically adjust evidence integration. We introduce a change-detection task in which both state and context must be inferred from a single stream of inputs in a dynamic and uncertain environment. Using this framework, we make three key contributions: (1) We show that mice rapidly adjust their behavioral strategy in response to implicit changes in input statistics, adapting within a single trial without reward or feedback. This suggests that their behavior is guided by a learned inference model rather than trial-and-error learning. (2) We derive the Bayes-optimal policy for the task, demonstrating that optimal inference requires a nontrivial real-time adaptation of evidence integration, in which both accumulation rates and decision boundaries adjust dynamically. (3) We show that artificial recurrent neural networks trained via reinforcement learning achieve near-optimal performance by embedding the sequential update of the Bayes-optimal decision variable within their recurrent dynamics. Together, our findings establish flexible perceptual inference as a fundamental computational principle, providing mechanistic insights into how biological and artificial systems adapt to uncertain and dynamically changing environments.

## 2 A change-detection task to study flexible perceptual inference

To investigate how agents infer both an evolving state and a shifting context from the same inputs, we designed a **change-detection task with variable sensory reliability**. The task requires tracking a latent state that determines reward availability while simultaneously adapting to changes in sensory reliability, which we consider the “context”. This simple dual-inference problem reflects the challenges of more general real-world problems.

### Task structure

The agent must infer a latent binary state, *s*_*t*_∈ {0, 1}, where *s*_*t*_ = 0 denotes an *unsafe* state and *s*_*t*_ = 1 a *safe* state. The state evolves stochastically: each trial begins in the unsafe state, transitioning to safe with probability *λ* per time step. The agent’s objective is to act (*a*_*t*_ = 1) only in a safe state to obtain a reward and to withhold action (*a*_*t*_ = 0) otherwise. A trial ends immediately upon action.

The agent does not directly observe *s*_*t*_ but instead receives a binary input, *x*_*t*_ ∈ {0, 1}, which we denote as *nogo* (*x*_*t*_ = 0) or *go* (*x*_*t*_ = 1). These observations are **noisy reflections of the current state**. A latent context parameter, *θ*_*t*_, determines sensory reliability: in an unsafe state, the input is veridical (*x*_*t*_ = 0) with probability 1 − *θ*_*t*_ and misleading (*x*_*t*_ = 1) with probability *θ*_*t*_. In a safe state, the observations are always reliable (*x*_*t*_ = 1) (Fig. 1a). The context parameter *θ*_*t*_ can change probabilistically between trials, grouping them into blocks of varying lengths (Fig. 1b).

**Figure 1:**
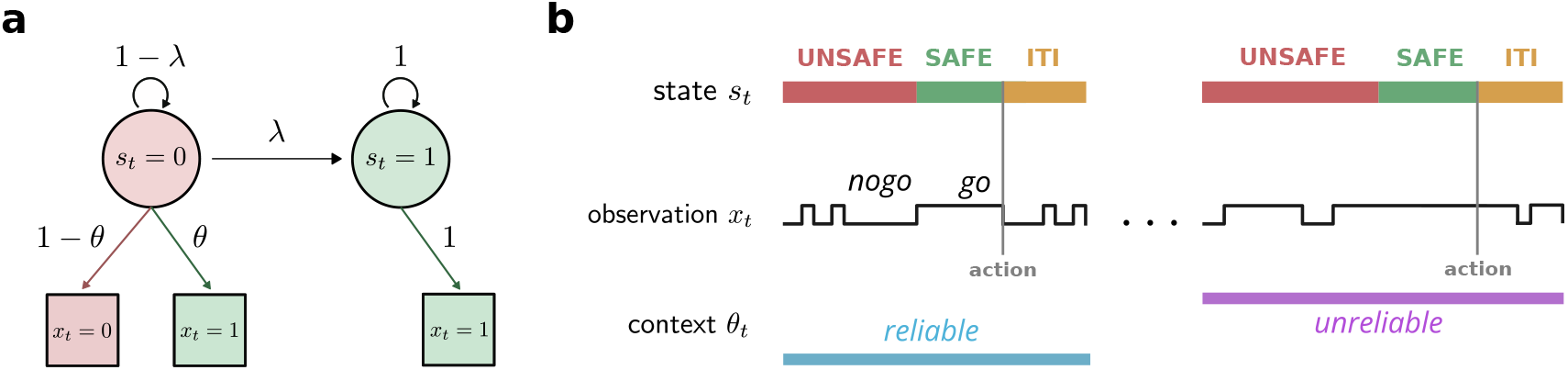
Task structure and trial dynamics. **(a)** Each trial follows a hidden Markov model with a binary latent state *s*_*t*_, which can be either *safe* (*s*_*t*_ = 1) or *unsafe* (*s*_*t*_ = 0). Trials begin in the unsafe state, and at each time step, the state transitions to safe with probability *λ*. During the unsafe state, the agent receives a *nogo* observation (*x*_*t*_ = 0) with probability 1 − *θ* or a misleading *go* observation (*x*_*t*_ = 1) with probability *θ*. The parameter *θ* defines the **context** and remains fixed throughout a trial. The agent earns a reward for acting only in a safe state, and each trial ends upon the first action. **(b)** Trials are separated by an inter-trial interval (ITI) and grouped into blocks with different values of *θ*, distinguishing **reliable** (low *θ*) from **unreliable** (high *θ*) contexts. The sequence of *go* and *nogo* observations provide noisy evidence about the hidden state and context. Following a block switch, agents are required to **simultaneously** infer both the state and the context.

### Optimal strategy and speed-accuracy trade-off

The agent faces a fundamental trade-off between acting prematurely (risking incorrect responses) and delaying too long (reducing the reward rate^10,42,43^). The optimal policy is characterized by a **waiting time**, defined as the number of consecutive *go* signals since the last *nogo*. This waiting time should depend on *θ*_*t*_: as sensory reliability decreases (higher *θ*_*t*_), the agent should wait longer to ensure that the state transition has occurred. Maximizing the reward rate over time thus imposes a **speed-accuracy trade-off**, balancing cautious waiting with efficient action. For each context *θ*, there exists a finite optimal waiting time *τ*^⋆^(*θ*) that maximizes the reward rate *r*(*τ*; *θ*).

### The challenge of ambiguous inputs

A key difficulty in this task is resolving input ambiguity. Let us consider a long string of consecutive *go* signals. This abnormal observation could indicate either that the state has transitioned to safe or that the context has shifted to a less reliable environment (*θ* increasing). To act optimally, the agent must **jointly infer the latent state and the context**—a form of flexible perceptual inference. Unlike traditional reinforcement learning, where adaptation is driven by trial-and-error, optimal behavior requires real-time inference within a single trial.

### Implications for flexible inference

To perform well, the agent must develop internal belief states that integrate incoming observations with prior expectations about state transitions and context shifts. In Section 3.1, we show that **mice trained on this task adapt their behavior within the first trial of a new context**, indicating that they infer context implicitly from observations rather than relying on reward feedback, motivating our study of flexible perceptual inference. In Section 3.2, we derive the **Bayesian-optimal solution** and analyze the dynamics of an ideal observer. Finally, in Section 3.4 and Section 3.6, we demonstrate that artificial neural networks trained on this task develop representations that mirror Bayesian inference.

## 3 Results

### 3.1 Trained mice adapt their behavior to a new context within the first trial

We trained adult mice in an instantiation of our task. Water-deprived, head-fixed mice were placed in front of a water delivery spout (Fig. 2a). In the experiments, we represented the *go* signals (*x*_*t*_ = 1) as a continuous auditory tone, while the *nogo* signals (*x*_*t*_ = 0) were represented by silence. The sound sequences were generated using the trial structure described in Section 2. Each signal lasted 0.2 seconds, with no gaps between consecutive signals. The mice were rewarded with a water drop for successful actions, defined as licking the water delivery port during the safe state. Detailed experimental procedures are provided in the STAR Methods.

**Figure 2:**
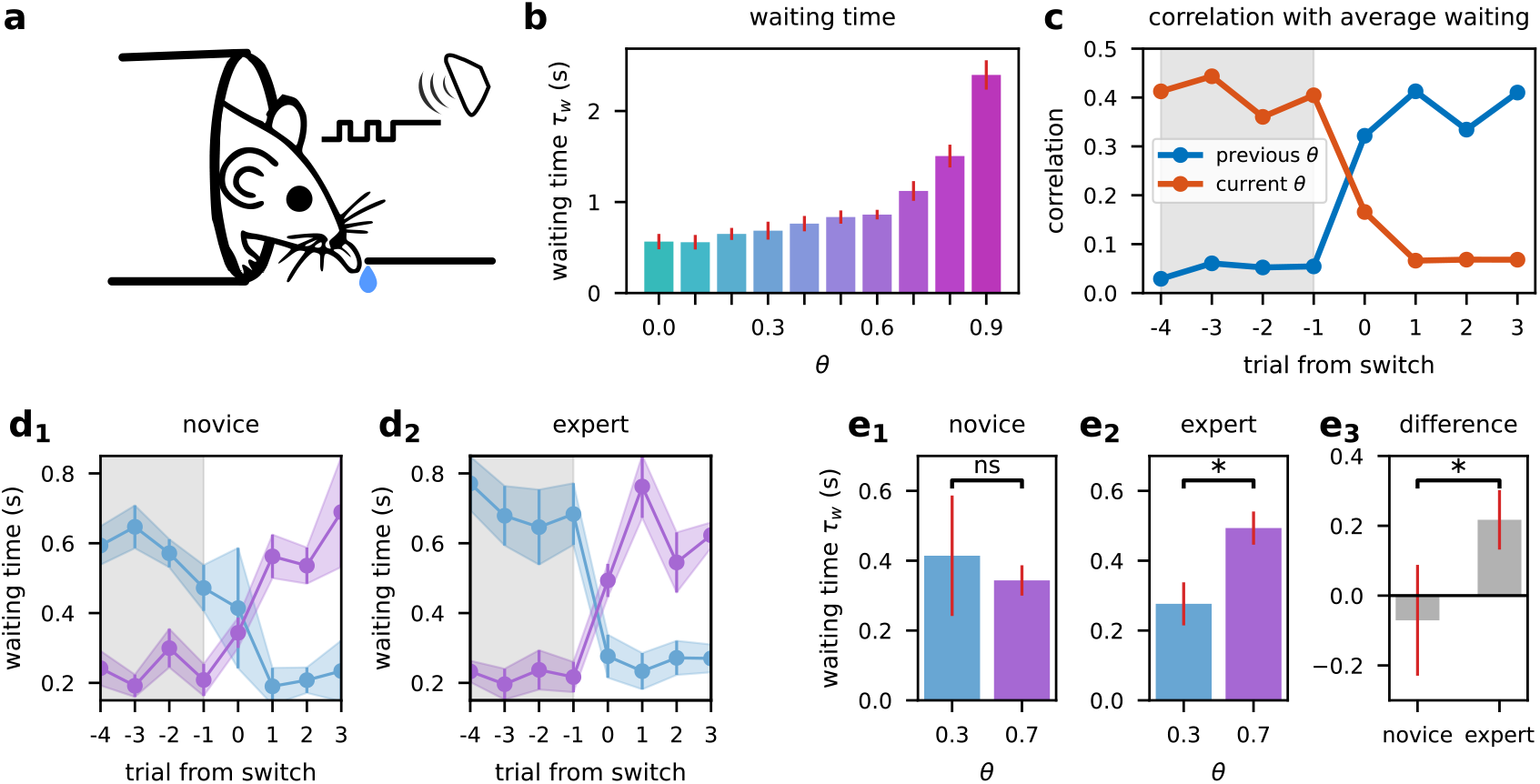
Mice adapt their waiting times to uncertainty without requiring explicit feedback. **(a)** Experimental setup. Head-fixed, water-restricted mice performed an auditory version of the task. Licking during the *safe* state resulted in water delivery, requiring mice to infer latent states over time. **(b)** Mean waiting time *τ*_*w*_ as a function of the contextual parameter *θ*. Mice exhibit longer waiting times in higher-uncertainty contexts. **(c)** Correlation between trial-by-trial waiting times and the average waiting times from the previous (orange) and current (blue) context blocks. Following a block switch (shaded region), mice rapidly adjust their waiting times to the new context, showing **first-trial adaptations** that rely on inference rather than feedback or reward. **(d)** Trial-by-trial waiting times aligned to block transitions in **novice** (first five sessions per mouse, left) and **expert** (last five sessions per mouse, right) animals. Data are shown separately for transitions from high to low uncertainty (*θ* = 0.7 →0.3, light blue) and low to high uncertainty (*θ* = 0.3 →0.7, purple). Experts exhibit more pronounced context-dependent waiting time modulation compared to novices. **(e)** Waiting times during the **first trial after a context switch**, summarizing the data from (d). **(e**_**1**_) Comparison between contexts in **novice** mice shows no significant difference. **(e**_**2**_**) Expert** mice exhibit significantly longer waiting times in higher-uncertainty contexts (*p <* 0.05, Wilcoxon signed-rank test). **(e**_**3**_) The difference in waiting time modulation between novice and expert mice demonstrates increased sensitivity to context with experience. Error bars and shaded regions represent standard error across mice.

To assess how mice adapt their behavior to different latent states of the environment, we trained and tested a group of 17 mice on our task, switching between blocks of varying values of *θ*. We focused on analyzing the mice’s *waiting times*, defined as the duration between the last *nogo* signal and the first lick. Since the underlying state is hidden, the number of consecutive *go* cues is hence the only relevant observable. In the next section, we derive the optimal solution in terms of waiting times.

Fig. 2b presents the average waiting times for different contexts, demonstrating that, consistent with normative predictions we detail in Section 3.2, mice wait longer as *θ* increases (Pearson’s r = 0.83, *p <* .005). Although the behavioral statistics show that mice modulate their waiting time given the latent state, they do not fully reveal what elicited this behavioral change. We were particularly interested in determining whether mice infer the hidden context from input statistics or adapt their behavior based on rewards (or lack thereof) following their actions. We reasoned that if mice relied solely on reward feedback, we would only observe a significant behavioral change from the second trial after the context had changed (i.e., after experiencing the first success or failure in the new context). In contrast, if mice had already adapted their behavior in the first trial, it would suggest that they were utilizing sensory-based inference to inform their actions.

### First trial adaptation

To investigate behavioral adaptation independent of feedback, we analyzed the mean waiting times in the first trials following each context change. Block switches occurred irregularly, preventing mice from anticipating context changes. In the first trial of a new context, mouse behavior already correlated more strongly with the steady-state waiting time of the new context than the old context’s (Fig. 2c). The significant change in behavior from the very first trial strongly indicates that the mice’s policy adapts to implicit changes in environmental conditions through inference rather than solely relying on behavioral feedback.

To demonstrate that the observed changes in waiting time represent learned task-related behavior, we conducted an additional experiment comparing mice at different stages of training. We first trained a new group of 7 mice on the task in a single context (*θ* = 0.3), allowing them to acquire basic familiarity with the environment. We then introduced these mice to the full task, which included unpredictable block switches between contexts with *θ* = 0.3 and *θ* = 0.7. This design allowed us to compare the performance of novice mice with that of mice with prolonged exposure to context switching.

Fig. 2d illustrates the mean response times, in novice (d_1_) and expert (**d**_**2**_) mice (see supplementary), for the last four trials before a block switch and the first four trials after the switch. Fig. 2e illustrates the mean waiting time for the first trial following a block switch (e_1_ and e_2_), and the behavioral difference of the first trial between contexts (e_3_). The results reveal a notable difference between the two groups. Compared to novice mice, expert mice demonstrated a significant change in waiting time from the very first trial of a new block.

This contrast between novice and expert performance indicates that the context adaptation we observe is indeed a learned task-related behavior. The ability of expert mice to rapidly adjust their waiting times in response to changes in context is not a simple response to auditory stimuli, nor is it merely a by-product of the increased frequency of *go* signals during high *θ* contexts. Instead, it represents a nontrivial cognitive skill acquired through experience with the task.

Our experiments with mice reveal three critical findings: (1) Mice adapt their behavior to the environment’s latent variables; (2) this adaptation is primarily driven by inference from auditory input statistics rather than reward feedback; and (3) the ability to adapt from the first trial is a learned behavior, developed through experience with changing contexts. These results demonstrate mice are capable of flexible perceptual inference, allowing them to rapidly adapt to dynamic environments without relying solely on feedback. In the following sections, we will develop a Bayesian theory for optimal latent state inference in reinforcement learning settings and explore potential neural mechanisms underlying this capability using artificial neural networks.

### 3.2 Optimal decision-making in a dynamic environment with latent interactions

The environment introduced in the previous section requires the agent to simultaneously track a latent binary state and a latent contextual parameter that governs observation reliability. These dual uncertainties give rise to a challenging inference problem. To illustrate, consider a sequence of *k* consecutive *go* signals ***x***_*t*−*k*:*t*_ = 1. In an *unsafe* state, such a sequence should prompt the agent to update its estimate of the context parameter *θ*, since repeated *go* observations in the unsafe state suggest a higher likelihood of observation flips. Conversely, in a *safe* state, the same stream of signals carries no information about *θ*. Further complicating matters, a transition from unsafe to safe can occur mid-sequence, confounding the inference of *θ* and demanding a simultaneous estimation of both the state and the context.

This complexity reflects a form of *negative interaction information* ^44,45^ between the latent state and the context: each new observation not only strengthens their statistical coupling, but also amplifies the uncertainty surrounding each individual variable. While the state and the context may be independent *a priori*, they become statistically dependent once conditioned on observations. Thus, resolving the joint inference of these hidden variables requires a unified treatment of their evolving likelihoods and the agent’s actions.

Identifying an optimal policy in this setting goes beyond mere inference, as it requires the agent to select actions that maximize rewards in the face of dynamic uncertainty. We thus employ the framework of partially observed Markov decision processes (POMDPs)^46^ to develop a rigorous Bayesian account of how an ideal observer should update its internal beliefs and choose actions. In the subsequent sections, we derive the belief-update equations for an optimal Bayesian observer and show how these updates drive a policy that achieves near-maximal reward. This analysis reveals how flexible perceptual inference can emerge through sequential belief-state updates that integrate observations with a learned internal model.

#### Markov approximation

The task in Section 2, designed to allow reliable mice behavior, contains two inherently non-Markovian features: (1) context changes occur only at trial boundaries, and (2) inter-trial intervals (ITIs) are sampled from a uniform distribution rather than an exponential distribution, which would satisfy the Markov property. To formulate the task as a partially observed Markov decision process (POMDP), we introduce two key approximations, both of which are well justified within the structure of our task.

First, we approximate context changes as a continuous-time Markov process. At each time step, the context can transition with a small probability *ϵ*, selecting uniformly from *m* discrete values 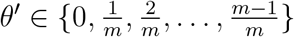.The context transitions are described by:

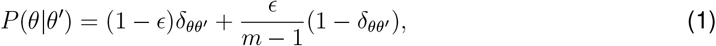

where 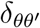 is the Kronecker delta function. This formulation approximates the discrete trial boundaries by allowing small, continuous changes in context over time. This approximation becomes exact as *ϵ* →0 and seamlessly generalizes to continuous value contexts as *m* → ∞. Thus, the choice of *ϵ* and *m* can be adjusted to balance computational feasibility with task fidelity.

Second, we treat ITIs as part of the *unsafe* state. This simplification is justified because ITIs share similar input statistics with the unsafe state, and well-trained Bayesian agents typically avoid taking actions during ITIs. By merging these intervals with the unsafe state, we reduce the system to two latent states: *safe* (*s* = 1) and *unsafe* (*s* = 0). The state transitions are then governed by:

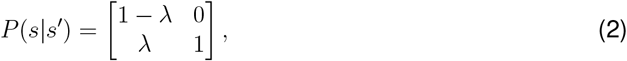

where *λ* is the probability of transitioning from the unsafe to the safe state.

By combining the independent transition functions in (2) and (1), the task can be approximated as a POMDP. In this framework, the agent observes only the binary input *x*_*t*_, while the two underlying latent variables, state *s* and context *θ*, evolve according to a Markov process. This representation retains the key features of the task while simplifying its mathematical formulation, enabling us to derive the optimal policy using the POMDP framework.

#### Belief states

In the framework of a POMDP, belief states provide a probabilistic representation of the agent’s knowledge about the environment, encapsulating all relevant information from past actions and observations^47^. Specifically, a belief state is a probability distribution over the latent variables, effectively transforming the POMDP into a fully observable Markov decision process (MDP) over the belief space. This transformation enables decision-making based on a sufficient statistic of the agent’s history, but solving the POMDP requires defining how beliefs are updated with new observations and how they guide optimal actions.

In our task, the belief state 𝒫_*t*_(*s, θ*|***x***_≤*t*_) represents the joint probability of the latent state *s*_*t*_ and the context *θ*_*t*_ given the observation history ***x***_≤*t*_. **Although** *s* **and** *θ* **are *a priori* independent, they become statistically dependent on observations**. This conditional dependence arises because the same observations inform both the current state and the context, creating intertwined estimates.

To illustrate this dependency, consider a sequence of *go* signals *x*_*t*_ = 1. As the sequence grows, the probability of being in the safe state increases, but it also increases the likelihood of a high-uncertainty context (larger *θ*). Resolving this interdependence requires maintaining and updating the full joint belief.

State and context estimators are derived from the posterior mean of the belief state:

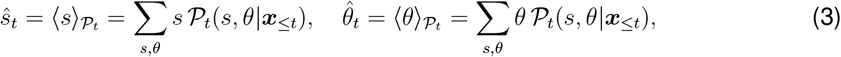

where 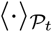 denotes averaging with respect to the current belief 𝒫_*t*_. These estimators, *ŝ*_*t*_ and 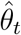,provide the agent’s best estimates of the state and context at any given time. The estimate evolves by updating the joint belief as new observations arrive.

#### Sequential updating of belief states

In dynamic environments, agents must continuously update their internal representations as new observations become available. In a Bayes-optimal setting, it is assumed that the agent has knowledge of the statistical structure of the environment, including the state transition probabilities *P* (*s*| *s*′), the context transition probabilities *P* (*θ*| *θ*′), and the observation likelihood *P* (*x*_*t*_ |*s, θ*). This knowledge could be acquired through prior experience or training. Using these known probabilities, the agent updates its belief state to optimally integrate new information with prior knowledge, ensuring that its internal belief reflects the current environment as accurately as possible.

We obtain a master equation for updating the belief state 𝒫_*t*_(*s, θ* |***x***_≤*t*_) using the Chapman-Kolmogorov equation^7,48^:

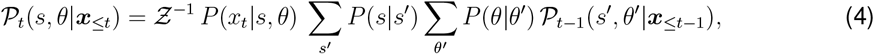

where Z is a normalization factor ensuring that 𝒫_*t*_(*s, θ*|***x***_≤*t*_) sums to 1.

On the left-hand side of Eq. (4), we have the current belief state. On the right-hand side, the update integrates the likelihood of the current observation *P* (*x*_*t*_|*s, θ*), which reflects how well the current input aligns with the latent variables, the transition probabilities *P* (*s*|*s*′) and *P* (*θ*|*θ*′) which describe the dynamics of state and context, and the prior belief state 𝒫_*t*−1_(*s*′, *θ*′| ***x***_≤*t*−1_) which incorporates all information available up to the previous time step. Together, these components enable the belief state to evolve as new information becomes available, combining prior knowledge with real-time observations. A detailed derivation of Eq. Eq. (4) can be found in the Supplementary Materials.

To initialize the belief state at the beginning of each trial, we assume that the world starts in the unsafe state (*s*_*t*_ = 0), leading to 𝒫_*t*_(*s* = 1, *θ*) = 0. The state in the previous step *s*_*t*−1_ is known because the agent has just acted and received feedback. Consequently, the initial distribution of *θ* is given by the belief state at the end of the previous trial, conditioned on the known state at that time: 𝒫_*t*_(*s* = 1, *θ*) = 0 and 𝒫_*t*_(*s* = 0, *θ*) = 𝒫_*t*−1_(*θ* |*s* = *s*_*t*−1_). This setup allows the agent to leverage feedback from the environment in addition to sensory observations.

#### Optimal policy

The optimal policy maximizes the expected reward rate, defined as the average reward over a fixed time horizon^10^. This requires balancing the need to gather sufficient evidence with the time cost of waiting, reducing the policy to determining the optimal number of consecutive *go* signals, *τ*, required before acting, which depends on the context *θ*.

For a given *θ*, the probability of being in the safe state after *τ* consecutive *go* signals is

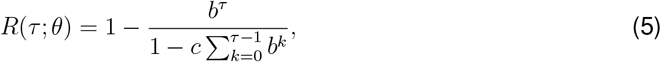

where *b* := (1− *λ*)*θ* is the probability of observing a misleading *go* signal, and *c* := (1 −*λ*)(1 −*θ*) is the probability of observing a *nogo* signal.

The expected trial duration is

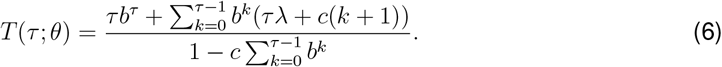

The reward rate is then given by

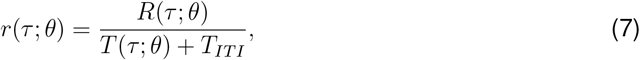

where *T*_*IT I*_ is the average inter-trial interval. The optimal waiting time *τ*^⋆^(*θ*) maximizes this reward rate

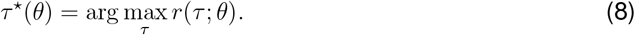

The curves in Fig. 3c illustrate the relationship between *r*(*τ*; *θ*) and *τ* for several values of *θ*. The maximum of each curve indicates the optimal waiting time and the corresponding reward rate for that specific context.

**Figure 3:**
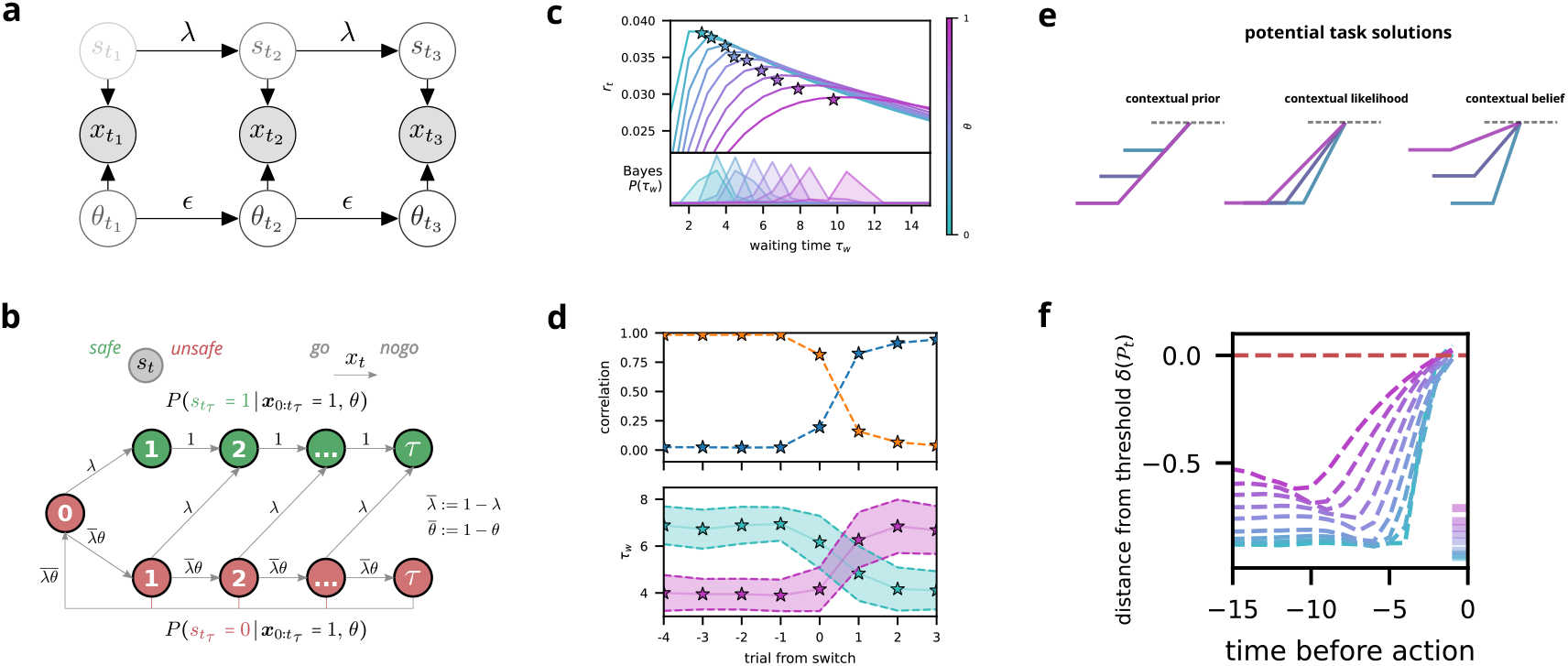
Optimal Bayesian inference of latent state and context. **(a)** Task structure modeled as a hidden Markov process. The latent state *s*_*t*_ and the context *θ*_*t*_ are independent sources of sensory observations *x*_*t*_, transitioning at rates *λ* and *ϵ*, respectively. Importantly, conditioned on the observations *x*_*t*_ the state and context are *dependent*. **(b)** Markov model describing the accumulation of *τ* consecutive *go* (*x*_*t*_) signals within a trial. Transitions between *unsafe* (red) and *safe* (green) states occur dynamically, with a *nogo* signal resetting the count *τ* to zero. This structure captures the probabilistic evolution of latent state estimates based on sequential evidence. **(c)** Top: Reward rate as a function of waiting time *τ*_*w*_ across different contexts, illustrating optimal action timing. A Bayes-optimal agent (stars) that simultaneously infers latent state and context achieves near-maximal reward rates. Bottom: Bayesian agent’s waiting time distributions across contexts. The variance is a result of uncertainty in the current context estimation. **(d)** Top: Correlation between trial waiting time and average waiting times from the previous (orange) and current (blue) blocks, demonstrating context-dependent, first-trial adaptation (cf. Fig. 2). Bottom: Average waiting times for specific context transitions (*θ*_*t*_ = 0.3 →0.7, purple; *θ*_*t*_ = 0.7 →0.3, light blue). Shaded areas indicate the standard deviation of the Bayesian posterior. **(e)** Alternative implementations of context-dependent decision models. Left: Contextual priors influencing baseline evidence. Middle: Contextual likelihood affecting evidence accumulation rate. Right: A full *contextual belief* model, where both baseline and accumulation dynamics are modulated by context, best matching Bayesian inference. **(f)** Trial-averaged trajectories of the Bayesian decision variable *δ*_*t*_, aligned to the time of action. The evolution of *δ*_*t*_ follows a structured ramping dynamic, with values during the last *nogo* trials (bottom right) varying systematically with context. The observed dynamics align with a **contextual belief** model, where both baseline levels and accumulation rates are shaped by the inferred context (cf. **d**, left).

In our task, the context *θ* is not directly observable and must be inferred from the input stream. The optimal waiting time, therefore, must account for this uncertainty by maximizing the expected reward rate, averaged over the belief state

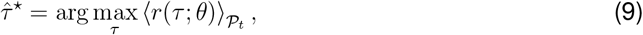

where the expectation is taken over the current belief 𝒫_*t*_(*θ*), reflecting the agent’s estimate of the context based on all past observations.

A key property of the task is that, following a *nogo* signal, the probability of being in a safe state drops to zero. This intuitive result arises directly from the task structure and can be formally derived from the dynamics in Eq. (4). Specifically, when a *nogo* signal is observed (*x*_*t*_ = 0), the likelihood of being in the safe state becomes zero, as *P* (*x*_*t*_ = 0|*s*_*t*_ = 1, *θ*) = 0. Conversely, with each consecutive *go* signal, the probability of being in the safe state 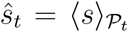 increases monotonically.

Using the optimal waiting time 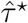, we calculate the belief threshold *ŝ*^⋆^ at which the agent should act. This threshold represents the minimum belief in the safe state required to maximize the reward rate:

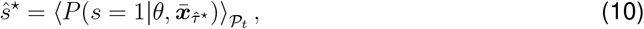

where 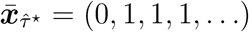 denotes a sequence starting with a *nogo* signal *x*_*t*_ = 0, followed by 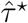 consecutive *go* signals *x*_*t*_ = 1.

This belief threshold *ŝ*^⋆^ defines the critical confidence level at which the agent should act. It captures the trade-off between collecting more evidence and the time penalty incurred by waiting too long. As the waiting time exceeds 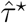,the expected reward rate decreases monotonically. Thus, the agent’s policy is to act whenever the current belief in the safe state *ŝ*_*t*_ reaches or exceeds the threshold *ŝ*^⋆^ ^49^.

Importantly, *ŝ*^⋆^ varies with the belief 𝒫_*t*_, and in particular with the estimated context 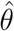.In more uncertain environments (higher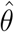), the threshold *ŝ*^⋆^ decreases, meaning the agent requires less certainty to act. This adjustment reflects the need to act more readily in contexts with ambiguous input to avoid excessive delays.

The belief threshold enables the formulation of a deterministic policy based solely on the current belief state

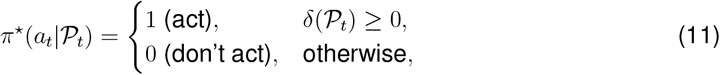

where we introduced the *decision variable δ*(𝒫_*t*_) = *ŝ*_*t*_ − *ŝ*^⋆^. This policy directs the agent to act when its belief in the safe state exceeds the optimal threshold, translating the agent’s understanding of state and context into a concrete action rule.

Together, the belief-update rule in (4) and the policy in (11) define a Bayes-optimal strategy for decision-making in dynamic environments. By inferring latent states and contextual reliability from a single observation stream, this framework integrates evidence in real time, resolves input ambiguity, and adapts to changing statistics; it provides a theoretical benchmark for understanding flexible perceptual inference.

### 3.3 Dynamics of Bayes-optimal perceptual decision-making in dynamic environments

Having derived a Bayes-optimal solution to account for the joint inference of state and context (Section 3.2), we now apply this theoretical framework to the change-detection task introduced in Section 2. The Bayesian solution allows us to quantify how an ideal observer integrates ambiguous observations and adapts to implicit context changes. Crucially, it also allows us to study how an ideal drift-diffusion model would implement simultaneous inference of the latent state and context.

#### Performance of a Bayes-optimal actor

We first assessed the performance of an agent that updates its belief states according to Eq. (4) and follows the optimal policy in Eq. (11). As illustrated in Fig. 3c (top), the Bayes-optimal actor attains near-perfect performance, closely approximating an agent with explicit knowledge of the context. The modest performance gap arises from residual uncertainty in the estimated context, which broadens the distribution of waiting times *P* (*τ*_*w*_) (bottom of Fig. 3c). This efficiency gap reflects the central challenge of *flexible perceptual inference*: using noisy and partial observations to infer, on the fly, both the binary state and the reliability of incoming signals.

#### Adapting to a new context within the first trial

One of the hallmarks of flexible perceptual inference, as laid out in the introduction, is rapid adaptation to changing environments without relying on trial-and-error. Our Bayesian framework meets this requirement by continuously updating the context estimate from each incoming observation. Fig. 3d demonstrates how the Bayes-optimal actor adapts within the very first trial of a new context block: correlations between waiting times in the current trial and in previous blocks reveal immediate behavioral shifts, even before any explicit feedback. This ability to adapt in real-time matches the behavior observed in mice (Fig. 2c,d), suggesting that animals may employ similarly sophisticated inference when faced with dynamic uncertainties.

#### Comparison with drift-diffusion dynamics

The drift-diffusion model (DDM) is a prominent framework for perceptual decision-making, effectively capturing continuous evidence accumulation and threshold-based decisions^10,11^. While earlier work has shown that DDMs implement near-optimal strategies for simpler value-based decisions^14^, it remains unclear how such models handle latent context shifts without explicit cues.

In our change detection task, the Bayes optimal strategy prescribes a context-dependent waiting time *τ*^*^(*θ*) that balances speed and accuracy under uncertainty. This dependence on *θ* can be cast in a DDM-like framework via two broad mechanisms:

1. **Contextual Prior:** Shifting the baseline of accumulation relative to the threshold so that higher *θ* (more unreliable inputs) places the baseline further from the threshold, prolonging the integration interval (Fig. 3e, left).
2. **Contextual Likelihood:** Altering the drift rate. Slower accumulation in high-*θ* contexts translates to longer waiting times, while faster accumulation in low-*θ* contexts yields more rapid decisions (Fig. 3e, center).

Although these two mechanisms are mathematically interchangeable under appropriate rescaling, practical implementations may yield distinct neural or behavioral signatures.

Interestingly, the exact sequential updates produced by Bayesian inference lead to **nontrivial decision-variable dynamics that can diverge from standard DDM assumptions**. In high uncertainty contexts (high *θ*), for example, the decision variable begins *closer* to the threshold rather than further away, as shown in Fig. 3f. This initial proximity is offset by a slower drift, ensuring that more *go* signals are needed to reach the threshold, thus delaying the action. For comparison, simpler DDM implementations that do not infer *θ* would not necessarily replicate such a combination of baseline shift and drift-rate modulation.

These distinctive dynamics reflect the underlying Bayesian logic of simultaneously inferring both state and context. In high-*θ* blocks, *go* signals are more likely to arise from false positives in the unsafe state, increasing overall uncertainty. This is captured by a lowered integration rate, where each *go* signal provides weaker evidence for the safe state. Meanwhile, the baseline remains higher because *go* signals are more frequent in high-*θ* contexts. **The combination of an elevated baseline and slower accumulation underscores the dual role of each *go* signal**: pushing the decision variable toward the threshold while signaling potential unreliability (Fig. 3e, right). Finally, the characteristic “dip” in the decision variable stems from the deterministic reset following the last *nogo* before action; the resulting change in baseline reveals the belief threshold *ŝ*^⋆^, which encodes the speed-accuracy tradeoff inherent in the task.

Our analysis shows that simultaneously inferring both state and context yields distinctive, contextdependent changes in the decision-variable dynamics that break the conventional logic of drift-diffusion models. These changes arise from the dual contribution of each observation, which informs both state and context estimates. In the next section, we demonstrate how recurrent neural networks trained via reinforcement learning spontaneously recapitulate these Bayesian-like updates, offering a biologically and computationally plausible mechanism for flexible perceptual inference.

### 3.4 Trained recurrent networks match Bayes-optimal performance

Equipped with the precise solution for the optimal behavior of a Bayesian agent and its unique dynamics, we now ask how neural networks solve the task. In particular, we show that a recurrent neural network trained using reinforcement learning solves the task optimally and that its dynamics align with the expectation of sequential Bayesian inference.

#### Network architecture and training

To solve our task, we used a deep reinforcement learning approach, implementing an actor-critic framework^50^ with 400 long short-term memory (LSTM) units^51^ (Fig. 4a). This architecture was chosen for its ability to handle sequential input in decision-making tasks and learn from temporal correlations in the input data. Although LSTM networks do not directly correspond to biological neurons, they allow us to study a neuronal implementation of the inference dynamics. Furthermore, LSTM networks have been suggested as models for the prefrontal cortex^52^ and can also be implemented using microscopic neuronal circuits^53^. Details on the network designs can be found in the supplementary Fig. S1.

**Figure 4:**
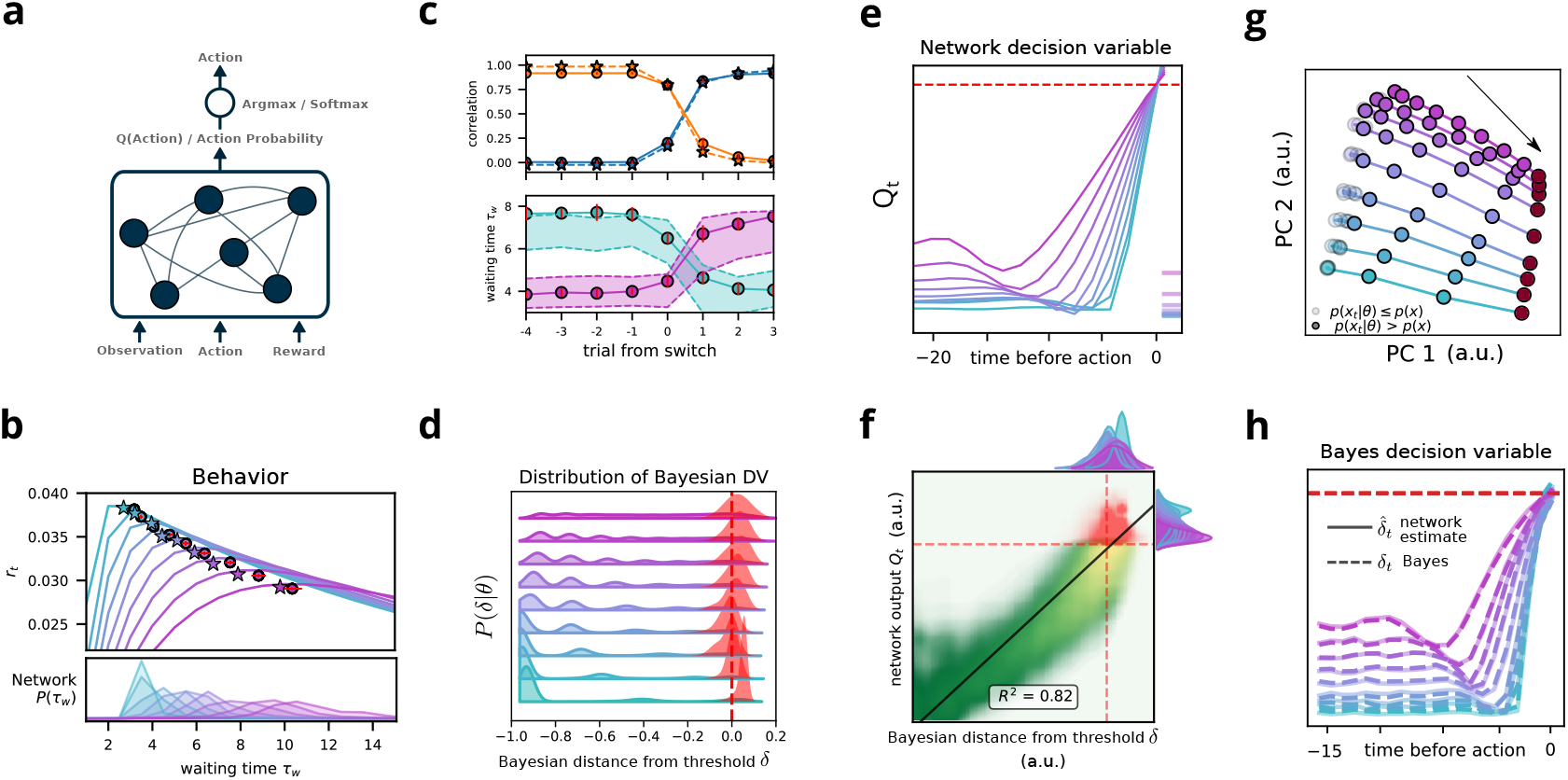
Neural networks trained via reinforcement learning approximate the sequential dynamics of the Bayes-optimal decision variable. **(a)** Schematic of the actor-critic LSTM network used for learning. The network receives observations, actions, and rewards as inputs and outputs action probabilities based on a learned decision variable *Q*_*t*_. **(b)** Top: Reward rate as a function of waiting time *τ*_*w*_ across different contexts. Trained networks (circles) infer both state and context, matching the performance of a Bayes-optimal agent (stars). Bottom: Bayesian agent’s waiting time distributions across contexts, providing an optimal benchmark for network behavior. Red error bars indicate standard deviation across networks (*n* = 10). **(c)** Top: Correlation of trial waiting time with the average waiting time in the previous (orange) and current (blue) blocks, showing first trial adaptation (cf. Fig. 2c, Fig. 3d top). Bottom: Mean waiting times for specific context transitions (*θ*_*t*_ = 0.3 → 0.7, purple; *θ*_*t*_ = 0.7 → 0.3, light blue), (cf. Fig. 2d and Fig. 3d bottom). Red error bars indicate standard deviation across networks (*n* = 10). **(d)** Distribution of the Bayesian decision variable during trials, grouped by context (color-coded curves), compared to its distribution at the moment the network acts (red). The alignment of action-triggered distributions with the Bayesian threshold suggests single-trial Bayes-optimal decisions. **(e)** Trial-averaged trajectories of the network’s decision variable *Q*_*t*_ aligned to the time of action. The temporal evolution of *Q*_*t*_ mirrors the Bayesian actor’s dynamics. The average value of *Q*_*t*_ during the last *nogo* trials (bottom right) is colored by context. **(f)** Scatter plot of network output *Q*_*t*_ versus the corresponding Bayesian decision variable *δ*_*t*_, showing a strong linear relationship (*R*^2^ = 0.82). Each point is colored by waiting time since the last *nogo* trial (*τ*, green to yellow) and at the moment of action (red), showing that network policies track Bayesian evidence accumulation throughout the trial. **(g)** Principal component analysis (PCA) of network dynamics. Trajectories are projected onto the first two principal components (PCs), revealing systematic structure in latent representations. Each marker represents a single time step, with color encoding context and transparency indicating temporal progression. **(h)** Time course of the decision variable *δ*_*t*_ extracted from network activity (solid lines) compared to the theoretical Bayesian decision variable (dashed lines). All data in this figure is averaged over 10 different trained networks (see STAR Methods for more details).

The critic subnetwork is designed to estimate the value function, predicting the current expected future reward. The output of the actor subnetwork, denoted *Q*_*t*_, parametrizes the probability of acting at each step, which is given by *P*_*train*_(*a*_*t*_ = 1) = (1 + tanh *Q*_*t*_)*/*2 during the training phase and *P*_*test*_(*a*_*t*_ = 1) = (1 + sign *Q*_*t*_)*/*2 during the testing. The sign function represents the deterministic threshold expected from our theory, while the soft threshold allows necessary exploration during training.

The network receives three inputs at each time step: the current observation, the action taken in the previous step, and the reward received in the previous step. This input structure allows the network to learn from the observations and previous actions and rewards. We trained the network using a reinforcement learning framework, specifically applying it to the task described in Section 2. Our training procedure used a Monte Carlo approach, which involves learning from complete episodes rather than after each time step or behavioral outcome^50^. After every 20 trials, each consisting of 30 steps on average, we update the network weights using backpropagation through time^54^.

#### Network performance

Our analysis demonstrates that trained RNNs closely approximate the performance and behavior of an optimal Bayesian agent across various context values (Fig. 4b, top panel). The RNNs achieve near-optimal performance, exhibiting qualitatively similar distributions of waiting times (Fig. 4b, bottom panels) compared to the Bayesian agents. Furthermore, networks display rapid context adaptation within the first test after a block switch (Fig. 4c), matching the behavior of an optimal Bayesian agent.

Despite the deterministic output of the network, response times varied across contexts. These variations in response times can be interpreted as environment-induced fluctuations in estimating the current context, akin to an optimal Bayesian agent.

### 3.5 Neural networks perform Bayes-optimal evidence accumulation

While our trained networks achieve Bayes-optimal performance on average, this observation alone does not demonstrate that they implement Bayesian inference mechanisms. To establish this stronger claim, we must examine the networks’ decision-making dynamics in individual trials. This presents a methodological challenge: We cannot directly compare the behavior of neural and Bayesian agents in parallel, as their actions actively shape the environment state. The independent operation of both agents would lead to divergent input histories, limiting us to comparing only their average performance.

To overcome this limitation, we developed an alternative approach: We allow the network to control the environment while computing the optimal Bayesian decision variable *δ*(𝒫_*t*_) based on the same input sequence. This analysis reveals that when the network initiates actions, the Bayesian decision variable consistently approaches its theoretical threshold, as evidenced by the distribution of *δ* at action times compared to its overall trial distribution in each context (Fig. 4d).

However, this threshold-crossing behavior alone does not conclusively demonstrate that the network implements Bayesian inference. Any evidence accumulation mechanism that integrates to a threshold would exhibit similar behavior. To establish whether the network truly captures the uncertainty inherent in Bayesian inference, we must examine its neural dynamics in detail.

A critical test of Bayesian computation lies in the network’s output dynamics during inference. When we examine the network’s decision variable *Q*_*t*_ aligned to action times across different contexts, we observe that the network reproduces the exact counterintuitive signature we derived for the optimal Bayesian solution (compare Fig. 4e with Fig. 3f). In contexts with higher uncertainty, where optimal integration requires longer waiting times, the network maintains its output closer to threshold during the final *nogo* signal while compensating with a slower integration rate.

This qualitative match with Bayesian dynamics is far from trivial. The network was only rewarded for implementing appropriate waiting times—it could have achieved this through simple context-dependent integration. Instead, it independently learned to adjust both the initial distance to threshold and the integration rate, precisely matching the optimal Bayesian solution. This strongly suggests that the network has discovered how to represent and account for the task’s inherent uncertainty, rather than merely learning a successful policy that approaches the Bayesian optimum on average.

To quantitatively validate the alignment between network computation and Bayesian inference, we examined the relationship between the network’s output and the Bayesian decision variable across entire trials. Our analysis reveals a strong correlation (*R*^2^ = 0.82) between these variables, even well before the decision threshold is reached (Fig. 4f). This correspondence demonstrates that the network has learned to track the Bayesian decision variable throughout the inference process, not just at decision time. Such continuous tracking explains why the network naturally exhibits the characteristic dynamical signature of optimal Bayesian computation.

#### Neural activity

Having established that the network’s output mirrors Bayesian dynamics, we next investigated how these computations are implemented in the network’s internal activity. Principal component analysis (PCA) reveals a remarkably low-dimensional solution: Three principal components (PCs) capture 88% of the total variance in neural activity. The first PC, accounting for 64% of variance, closely tracks the network’s action values *Q*_*t*_ (Fig. 4g), providing a direct neural representation of the decision process. The second PC (13.4% variance) encodes context information, while the third PC (10.6% variance) corresponds to post-action reset dynamics, ensuring proper initialization for subsequent trials (see STAR methods and Supplementary Materials for further details and the full PCA analysis).

Although this geometric description suggests a structured internal representation, it alone cannot verify *how* the network computes the Bayesian output. To establish this connection directly, we examined whether the network activity explicitly represents the key variables of our Bayesian theory. We constructed linear decoders with time- and context-independent readout weights to extract both latent variables from neural activity independently.

The decoding results provide compelling evidence for explicit Bayesian computation: Using 10-fold cross-validation with an 80-20 train-test split, we achieved near-perfect reconstruction of the Bayesian decision variable (*R*^2^ = 0.993, Fig. 4h) averaged across network instantiations. For comparison, decoding accuracy for the ground-truth state (logistic regression, accuracy= 0.9) and context (linear regression, *R*^2^ = 0.92) were also remarkably high. These results demonstrate that the network not only reproduces Bayesian dynamics in its output but also maintains explicit representations of the key computational variables in its neural activity.

Our analyses reveal that neural networks learn to approximate the sequential dynamics of the Bayesian decision variable without necessarily implementing a full Bayesian computation. This distinction is crucial—while Bayesian inference requires maintaining and updating joint probability distributions over latent variables, the network has discovered an efficient solution that captures the essential dynamics of the decision variable updates. The network could have adopted simpler strategies, such as basic evidence accumulation with context-dependent thresholds. Instead, it discovered how to approximately mirror the complex, context-dependent dynamics of optimal sequential inference, including precisely calibrated integration rates and initial conditions. This emergence of sophisticated computational strategies, derived purely from reward signals, reveals how reinforcement learning can give rise to principled solutions in dynamic inference tasks without explicit instruction in probabilistic computation.

### 3.6 Sequential Bayesian joint inference is implemented in the recurrent dynamics

A recent study showed that optimal Bayesian estimators can emerge from neural networks trained for value prediction^55^. In particular, estimators over beliefs can be linearly decoded even from untrained networks, suggesting that the intrinsic dynamics of random networks might be sufficiently rich to support Bayesian computation. In this case, Bayesian inference can be learned through reservoir computing^56^, training only the readout weights.

However, there are key differences between value prediction and perceptual decision-making. In particular, we consider an *operant* task that demands discrete decisions. The key distinction lies in the behavioral output. Whereas value prediction tasks involve estimating a continuous variable, our task requires categorical choice (act or wait) based on a dynamic decision variable. This difference is crucial, as optimal performance in our task depends on accurate single-trial state estimation. The average network output, which might suffice for value prediction, may not capture the temporal precision required for adaptive behavior in dynamic environments.

To examine whether random networks can maintain belief-like representations in this more challenging task setting, we used a “naive” model. This model consists of an RNN with fixed random weights for both recurrent and input connections. As in reservoir computing, only readout weights were trained, which feed into a *softmax* layer to produce action probabilities(Fig. 5d).

**Figure 5:**
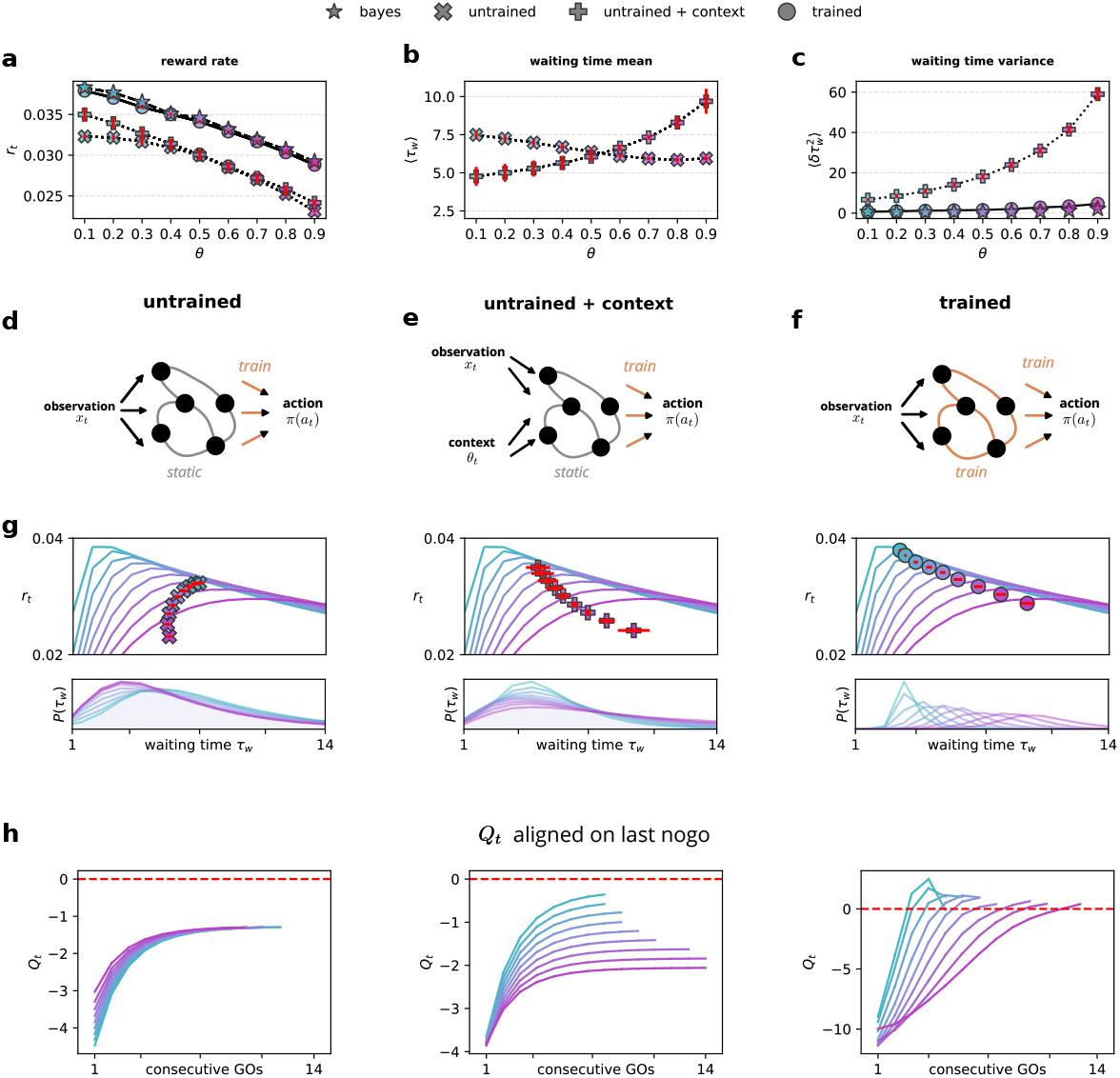
Learning recurrent dynamics is necessary for implementing sequential inference of both state and context. **(a)** Reward rate as a function of the context parameter *θ* for different network models (symbols indicated in the legend). Only the trained networks achieve Bayesian-level performance. **(b)** Mean waiting time ⟨*τ*_*w*_⟩ as a function of *θ* for untrained (×) and untrained + context (+) networks. The inclusion of an explicit external context improves the ability of a linear readout to modulate waiting times, increasing them for higher *θ*. **(c)** Variance of waiting times 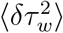 as a function of *θ* across models Bayes-optimal behavior, as well as trained and untrained + context networks. The high trial-to-trial variability of the untrained models contributes to their reduced performance. **(d–f)** Network schematics illustrating the different models. **(d)** Untrained networks receive only raw observations *x*_*t*_ but lack learned dynamics. **(e)** Untrained + context networks receive explicit context information *θ*_*t*_ alongside observations. **(f)** Trained networks learn recurrent dynamics that implement the sequential dynamics of the Bayes-optimal decision variable. **(g)** Top: Relationship between waiting times *τ*_*w*_ and average reward rate *r*_*t*_ across models, compared to the context-aware Bayesian reward curve (see Figs. 3 and 4). Bottom: Distributions of waiting times for different models, illustrating the effect of learning on action timing. **(h)** Evolution of the decision variable *Q*_*t*_ as a function of the number of consecutive **Go** (*go*) signals across different contexts. Left: Untrained networks fail to distinguish contexts, relying on stochastic behavior. Center: Providing explicit context information enables some degree of separation in decision dynamics. Right: Trained networks implement an optimal deterministic policy, rapidly adapting *Q*_*t*_ to the appropriate context.

We found that retaining the *softmax*-based probabilistic action selection was essential for the functionality of the naive model. This differs from the fully trained network, which could learn a deterministic policy as prescribed by the theory (using a hard threshold in the final layer). However, the naive model required stochastic action selection to perform acceptably.

The performance of this random RNN model was poor (Fig. 5a). Furthermore, it exhibits a different qualitative behavior. While optimal performance requires longer waiting times at higher *θ* values, the random RNN model showed the opposite pattern, with decreased waiting times at higher *θ* values. (Fig. 5b). This maladaptive behavior is a direct result of stimulus statistics. Without proper context inference, the network responds to go cues by simply increasing its action probability, ignoring the broader temporal structure of the task.

Despite the poor behavioral performance, we could decode Bayesian state and context estimators from the network’s activity with relatively high fidelity. Using the same cross-validation methods applied to trained networks, we found that random dynamics had strong predictive power, albeit slightly lower than fully trained networks (*p*(correct) = 0.89 ± 0.001 for the state and *R*^2^ = 0.91 ± 0.003 for the context estimators; mean ±std). These results align with the findings of^55^, which successfully decoded Bayesian estimators from random networks in a value prediction task. However, they raise the question of why the network’s performance still is so poor.

### Untrained networks fail when it matters the most

The apparent contradiction between the successful decoding of Bayesian estimators and poor behavioral performance (Fig. 5g left) in random RNN models stems from a specific limitation in their representation of latent variables. While network activity allows good decoding on average, it fails critically at the most behaviorally relevant moment, after several consecutive *go* signals, when the agent needs to make a decision.

Specifically, when the decision variable approaches the action threshold, the random network output *Q*_*t*_ shows no meaningful differentiation between contexts or the number of consecutive *go* signals (Fig. 5h left).

Random recurrent networks have inherent limitations in their temporal integration capabilities. Classical random networks exhibit exponentially decaying memory traces^57^, while random LSTM architectures can maintain somewhat longer dependencies^58^. This memory constraint allows linear decoders to differentiate contexts based on recent history, but the discrimination inevitably fails after several consecutive go signals as the memory trace degrades. Assuming a random network, the dynamics of the output *Q*_*t*_ can be captured by a simple linear integration model where

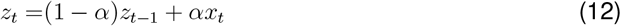

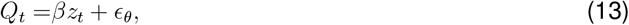

*z*_*t*_ represents a leaky linear integration of the observations *x*_*t*_ and *α* controls the memory trace. The output *Q*_*t*_ is a linear transformation of the integrator state with gain *β* and baseline *η*_0_. This minimal model accounts for 99.0 ± 0.2% of the single-trial variance while explaining only 74.0± 1.6% of the variance in fully trained networks, confirming that random networks implement a fundamentally simpler computational strategy.

### Learning the recurrent weights is required even when explicit context is provided

The failure of random networks to maintain context-dependent representations near decision boundaries raises a fundamental question: Does this limitation stem from an inability to infer context or from an inability to implement proper context-dependent integration even with perfect context knowledge? To differentiate between these possibilities, we modified our random RNN model by directly supplying the context as an additional input (Fig. 5e) (see Section 4 for technical details). This approach isolates the role of context inference from the computational demands of context-dependent evidence accumulation.

With explicit context information, the random network model demonstrated a significant qualitative improvement in its behavioral strategy. In particular, the mean waiting time increases with *θ*, as expected from optimal behavior and observed in mouse data (Fig. 5b). This qualitative change suggests that access to context information enables the network to implement a basic form of context-dependent integration. Still, quantitative analysis revealed persistent suboptimal performance (Fig. 5g middle), particularly pronounced at high *θ* values where reward rates remained significantly below those achieved by the Bayesian model and fully trained networks (Fig. 5a).

The random RNN model shows decreased performance despite achieving appropriate mean waiting times due to excessive response variability (Fig. 5c). This limitation comes from the fixed memory capacity of the network, which restricts its ability to integrate *go* cues beyond a certain duration reliably. To compensate for limited memory and achieve longer mean waiting times, the network implements a suboptimal strategy: maintaining the output *Q*_*t*_ at a context-dependent distance from the stochastic threshold, effectively trading off precision for approximate temporal control. This bias-variance trade-off allows the network to increase average waiting times through reduced action probabilities but at the cost of high trial-to-trial variability.

Similarly to the random network that did not receive an explicit context input, this mode can be explained by the linear integrator model *z*_*t*_ shown in (12), where the readout is given by *Q*_*t*_ = *βz*_*t*_ + *ϵ*_*θ*_. Here, the baseline depends on the explicit context through the set of parameters *ϵ*_*θ*_. As before, the linear model explains the output trajectory of the random model (with explicit context) very well (explained variance 99.1 ± 0.6%) while failing to explain fully trained networks (74.0 ± 1.6%) (Fig. 5h).

This systematic analysis of random networks reveals fundamental limitations in implementing Bayes-optimal behavior, even when provided with perfect context information. The now necessary reliance on stochastic policies starkly contrasts the deterministic Bayesian solution that uses internal belief states. Linear modeling exposed specific computational deficits: Random networks do not modulate their integration rate or decision threshold. When context is provided, random networks improve their behavior by making crude threshold adjustments. In contrast, trained recurrent networks (Fig. 5f) develop internal dynamics that independently control both integration rate and baseline in the precise and complementary manner predicted by Bayesian theory. These results demonstrate that optimal inference in our task emerges specifically through the training of recurrent connectivity, supporting a mechanistic link between reinforcement learning and flexible perceptual inference.

## Discussion

Our study explores adaptive decision-making in partially observed dynamic environments using a change detection task that requires the integration of evidence. This task involves two latent states that, while independent in isolation, become dependent when conditioned on past observations. We show that mice adapt their behavior within the first trial of a context change, relying on sensory inference rather than feedback. This behavior demonstrates flexible perceptual inference, enabling the disambiguation of observations and dynamic adjustment to implicit contextual changes. To uncover the computational principles behind flexible perceptual inference, we derived the Bayes-optimal strategy for reward rate maximization. The task’s conditional dependencies necessitate updating the joint distribution of latent variables. The optimal strategy manifests itself as an integration model in which both the accumulation rates and the bounds dynamically adjust in real time, reflecting the rapid adaptation to implicit changes in context. This flexibility in evidence integration results in counterintuitive dynamics of the Bayesian decision variable: In contexts with higher uncertainty, the decision variable is set closer to the decision threshold, offset by slower evidence accumulation.

We then trained recurrent neural networks (RNNs) on the task using reinforcement learning. The RNNs, while not explicitly constrained to Bayesian computations, achieved near-optimal performance. Their outputs mirrored the nontrivial dynamics of the Bayesian decision variable, adapting integration rates and thresholds to real-time contextual changes. This adaptation was enabled by learned recurrent connectivity, underscoring the capacity of neural networks to implement complex, context-dependent inference strategies based solely on reward-based training.

### Inference-based learning and cognitive flexibility

Our findings strengthen the growing evidence for model-based reasoning in animals^59–61^, providing a concrete neural example of this capability. We interpret dynamic adjustments in evidence integration rates and decision bounds as hallmarks of an adaptive internal inference model that supports flexible perceptual inference. This ability to update internal representations based on sensory input statistics, rather than relying solely on reward feedback, suggests a form of inference that transcends simple stimulus-response associations or purely reward-driven learning. From a computational perspective, our work extends the framework of meta-learning^50^—the concept of *learning to learn*—to encompass *learning to infer*. By demonstrating that recurrent neural networks can acquire near-optimal Bayesian inference through reinforcement learning, we propose a plausible neural mechanism underlying the cognitive flexibility exhibited by animals in dynamic environments.

### Counterintuitive dynamics as a signature of Bayesian inference

While extensive research has shown that animals integrate various sources of uncertainty into their decision-making^2,61–65^ and has linked computational models of perceptual decision-making to artificial neural networks^55,66,67^, identifying direct neural correlates of *Bayesian* decision-making remains challenging^68–70^. Our findings reveal a distinctive hallmark of fully Bayesian inference in dynamic, uncertain contexts: in blocks characterized by greater input unreliability, the decision variable starts *closer* to the threshold but accumulates evidence *more slowly*, reflecting the fact that each observation informs *both* the state and the context. This behavior stands in contrast to conventional drift-diffusion models, which often encode uncertainty solely via adjustments in drift or threshold distance^13^. Notably, we show that artificial neural networks trained in our task recapitulate this counterintuitive dynamic, indicating that joint inference over context and state can emerge naturally under reinforcement learning. This dynamic signature may serve as a valuable indicator of flexible, model-based computation in neural circuits, particularly in regions like the orbitofrontal cortex—an area implicated in planning and rapid adaptation^60^—where the neural underpinnings of Bayesian inference can be further explored.

### Recurrent neural networks as models for complex cognitive processes

Training RNNs to develop task-specific dynamics is essential for neural modeling of cognitive processes in complex, uncertain environments. We found that relying solely on readout weights from random RNNs failed to capture the intricate computations required for our task, consistent with the limitations of reservoir computing in temporally dependent tasks^56^. In contrast, training the recurrent connectivity enabled the networks to perform probabilistic computations in partially observable environments, where both latent states and contextual information must be inferred simultaneously. These results extend previous work demonstrating the role of recurrent dynamics in probabilistic computations^67,71,72^ to more complex cognitive tasks. By training RNNs on such tasks, we uncover neural mechanisms underlying flexible perceptual inference. This approach bridges theoretical models and experimental neuroscience, offering a powerful framework for studying the mechanisms underlying complex cognitive functions.

### Importance of operant tasks in studying decision-making

Our findings highlight the critical role of operant tasks in studying decision-making, as they reveal behaviors that cannot be captured by state estimation or reward prediction alone. While latent variables and their Bayesian expectations can be decoded even from random networks^55^, such networks exhibit poor task performance. This demonstrates that merely reading out mean or typical values from a network, as often done in reinforcement learning studies, is insufficient to assess its computational principles. By examining single-trial behavior in operant tasks, we combine normative principles with network simulations, revealing the subtleties of learned neural representations. This approach highlights the need for caution when interpreting neural data based solely on decoding specific variables, as it may obscure the underlying computational principles^73^.

In summary, our results establish that the rapid trial-by-trial adaptation seen in both mice and artificial neural networks arises from a unifying principle of *flexible perceptual inference*, whereby latent states and contexts are jointly inferred. By demonstrating how recurrent connectivity can naturally implement Bayesian-like computations under reinforcement learning, we highlight a plausible computational basis for real-time cognitive flexibility. These findings provide a computational and mechanistic foundation for real-time cognitive flexibility, offering testable predictions for neural circuits engaged in adaptive decision-making, for example, in the orbitofrontal cortex. Future work should aim to identify how such inference is implemented at the circuit level and how it interacts with reinforcement learning to shape behavior in dynamic environments.

## RESOURCE AVAILABILITY

### Lead contact

Requests for further information and resources should be directed to and will be fulfilled by the lead contact, Jonathan Kadmon.

### Data and code availability

We provide the code to our simulations at https://github.com/kadmon-lab/flexible-perceptual-inference.

## ACKNOWLEDGMENTS

This work was supported by the Gatsby Charitable Foundation, the National Institute of Psychobiology (J.K.) and the Israel Science Foundation (grant #1269/20 to E.L.). We thank Alkesh Yadav, Vladimir Shaidurov, and Lior Fox for their helpful discussions. E.L. is the incumbent of the Sachs Family Faculty Development Chair in Brain Sciences.

## AUTHOR CONTRIBUTIONS

J.S., J.B., and J.K. conceptualized the work with contributions from E.L.; J.S. developed the simulations and analyzed simulation data, with contributions by J.B.; J.B. and J.S. developed the theoretical results. H.R. and G.M.. conducted the animal experiments with contributions by J.S.; J.S., H.R. and G.M. analyzed the animal data; J.S., J.B., and J.K. wrote the manuscript, with contributions from H.R., G.M., and E.L; J.K. and E.L supervised the work.

## DECLARATION OF GENERATIVE AI AND AI-ASSISTED TECH-NOLOGIES

During the preparation of this work, the author(s) used OpenAI ChatGPT in order to edit and revise portions of text. After using this tool or service, the author(s) reviewed and edited the content as needed and take(s) full responsibility for the content of the publication.

## STAR METHODS

### Animals

All experiments were conducted using male C57BL/6 mice under water restriction (age: 2-4 months, weight: 20-30 g). Mouse weight was monitored daily and extra water was provided if needed to ensure that mice maintain no less than 80% of their original weight. Mice were housed in a 12-hour light/dark cycle with ad libitum access to food. All procedures were approved by the Institutional Animal Care and Use Committee (IACUC) of Hebrew University.

### Surgical procedures

To implant head bars for head fixation, mice were anesthetized using isoflurane (SomnoSuite, Kent Scientific USA) and placed in a stereotaxic apparatus (KOPF 962, David Kopf instruments, USA). Meloxicam (2 mg/kg) was injected subcutaneously for systemic analgesia and 0.1 ml lidocaine (2% solution) was injected subcutaneously before incising the scalp and exposing the skull. A 20 mm long and 3 mm wide head-post was then cemented to the skull horizontally using dental cement (C&B Super-bond dental cement, Sun Medical, Japan) and secured using dental resin (Pi-ku-plast, Bredent, Germany). Post-operative analgesia was administered and mice were allowed to recover for 2 to 3 days before beginning a 1-week water restriction schedule prior to training.

### Animal task implementation

Head-fixed mice were trained to perform an auditory change detection task. Each trial was discretized into 0.2-second time bins, during which the trial remained in one of four latent states: *unsafe, safe, premature*, or inter-trial interval (*ITI*). In all states, except the safe state, beeps (i.e., go cues) were generated with probability *θ* (meaning that if *θ* = 0.3, a beep occurred in 30% of bins). In the *safe* state, beeps occurred with probability 1 (i.e., in every time bin).

Each trial began in the *unsafe* state. If the mouse withheld licking, the state transitioned from *unsafe* to *safe* after a delay of 0.8 + 0.2 ⌊*x*⌋ seconds (where *x*~ *exp*(*λ* = 0.1) and ⌊*x*⌋ is the floor of *x*). Licking prematurely during the *unsafe* state triggered the *premature* state, which lasted for 7 seconds, before the state transitioned to the *ITI*. In the *safe* state (which could last up to 7 seconds), licking was rewarded with a 4–6 *µL* drop of water, and caused an immediate transition to *ITI*. The *ITI* duration was uniformly distributed between 1 and 5 seconds, and afterwards the next trial began.

Beeps consisted of a harmonic chord composed of pure tones at 2, 4, 6, 8, and 16 kHz, presented via a speaker adjacent to the mouse at 70 dB SPL. The task was controlled using the Bpod behavior measurement and control system (Sanworks, USA). Licking behavior was monitored using a 200 fps camera (Blackfly BFS-U3-16S2M-CS, FLIR, USA) with a Tamron M118FM16 lens (Tamron, Japan). Licking events were detected online using Bonsai^74^.

### Animal training

Two batches of mice were trained: the first batch (17 mice) was trained on all tested *θ*-values (0.1-0.9), while the second batch (7 mice) was trained on *θ* = 0.3 and 0.7 only.

For the first batch, initial training was conducted using a single *θ* value, with the first session using *θ* = 0.2 or 0.3, followed by 3–9 sessions in which *θ* values ranged from 0.1 to 0.7. Once mice demonstrated task acquisition by reliably responding to the state change, sessions with multiple *θ* values were introduced, presented in blocks of 30–100 trials. Mice were considered experts upon reaching stable performance, which occurred after ~ 10 training sessions. Upon achieving expert-level performance, we collected 10 sessions per mouse for analysis (Fig. 2 b, c).

The second batch (Fig. 2d,e) underwent 5 initial training sessions with *θ* = 0.3, followed by sessions with both *θ* = 0.3 and 0.7 in blocks of 50–100 trials. The first 5 sessions with both contexts (sessions 6, 7, 8, 9, and 11; session 10 was excluded as it only included *θ* = 0.3) were classified as ‘novice’ data. The last 5 training sessions (sessions 15–19) included *θ* = 0.3 and 0.7 in blocks of 30–60 trials and were classified as ‘expert’ data.

### Animal behavior analysis

Behavioral data were analyzed in MATLAB (MathWorks, USA). Trials were included in the analysis if the mouse licked at least once and if the trial occurred within the first 300 trials of the session. The first four trials of each session were excluded from adaptation analyses (Fig. 2c–e) because no prior context was available in the first block for mice to adapt their behavior relative to a preceding condition.

For correlation analyses, block averages were computed using all trials in a block except the first and last four trials. To examine the relationship between waiting time and block transitions (Fig. 2c), we calculated the Pearson correlation between waiting times of individual trials and the mean waiting time of the preceding or current block.

Waiting time differences (Fig. 2d) were computed as the average first-trial waiting time in *θ* = 0.7 minus the average first-trial waiting time in *θ* = 0.3. Two-sided Wilcoxon signed-rank tests were used to compare first-trial waiting times (N = 7 mice) (Fig. 2e). Pearson’s correlation coefficients and significance values were computed using MATLAB’s corrcoef function. Error bars indicate standard error across mice.

### Neural network training

**Architecture** We used the PyTorch library to implement an Actor-Critic framework^75^. Both the Actor and the Critic consisted of an LSTM layer (100 hidden units) followed by a linear readout: the critic readout produced a scalar value (*V*), while the actor readouts produced *Q*(action) and *Q*(inaction), passed through a softmax to yield action probabilities. We defined *Q*_*t*_ = *Q*(action) − *Q*(inaction). For standard policy execution, we sampled from the softmax distribution; when comparing to the optimal Bayesian agent, actions were chosen by argmax.

### Input and initialization

At each time step, both LSTMs received four inputs: (i) a scalar representing the current go/no-go state, (ii) the previous action, (iii) the previous reward, and (iv) optionally, the scalar context *θ*_*t*_ (for the untrained + context variant). Weight parameters were initialized using PyTorch’s default linear layer initialization, and LSTM hidden states were zero-initialized before the first episode. For subsequent episodes, hidden states were carried over from the final time step of the preceding episode to avoid signaling the start of a new episode.

### Training procedure

Networks were trained for 10,000 episodes, each comprising 20 trials under a single *θ* (sampled from 20 equally spaced values in [0, 1]). Each trial ended when the network responded. We applied a policy-gradient update with an entropy bonus of 0.05 to the Actor, while the Critic loss was scaled by 0.01 relative to the Actor^50^. Critic weights were updated by minimizing 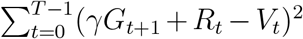, with a discount factor *γ* = 0.5. The final return *G*_*T*_ was set equal to *V*_*T*_. Optimization used RMSprop with an initial learning rate of 0.0005, decaying by 0.99 every 10 episodes, and momentum/weight decay both set to 1e-3. Gradients were computed via backpropagation through time at the end of each episode, then detached.

### Task and reward Structure

The *unsafe* state duration was drawn from an exponential distribution with a mean 10 steps (minimum of 1), always beginning with a forced no-go step. The intertrial interval (ITI) was uniformly distributed between 5 and 25 steps. **During training**, correct actions in the *safe* state delivered a reward of +1, whereas acting during the stimulus period incurred a negative reward penalty (*r* = − 3) to stabilize learning. At inference time, this negative reward was removed.

### Evaluation

We trained 10 networks per model variant (trained RNN, untrained RNN, and untrained RNN + context). Variability across the 10 networks could result from stochasticity in the weight-initialization, state durations, stimuli, and policy. After reaching comparable performance across all networks, the weights were frozen, and 5,000 episodes per network were evaluated and pooled for the final behavioral analysis.

## Analysis of network data

### Principle component analysis

PCA was performed on the Short-Term Memory stream of the Actor LSTM. Data from all units across all trails (including intertrial intervals) were centrelized and used to compute the principal components using Pythonn’s *scikit-learn* PCA package.

### Fitting network output to the linear intgrator model

To model the output of the recurrent network over time *Q*_*t*_ as a linear integrator, we optimize the model parameters under constraints 0 *< α <* 1, *β >* 0, and *ϵ*_0_ *>* 0. Optimization was performed using the first 100,000 data time steps, while the entire dataset was used to evaluate goodness of fit (Fig. 4g,h). The variability between the network realizations (*n* = 10) was negligible and did not affect the estimated fit parameters.

### Regression from Network Activity

To decode ground truth and Bayesian estimates, we performed regression on the short-term memory stream of the Actor network, as it represents neural activations relevant to decision-making. The Critic network, while essential for learning, does not contribute significantly during inference and was therefore excluded^50^.

All regressions were performed on trial data, excluding inter-trial intervals (ITI), as they are not behaviorally relevant. To obtain a linear readout, we applied logistic regression to decode the true state *s*_*t*_ and the Bayes optimal state estimate *ŝ*_*t*_, with the latter regressing to its log-likelihood ratio:

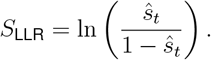

Context decoding was performed using linear regression to predict the true context *θ*_*t*_ and the Bayesian estimate 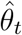.To decode the Bayesian decision variable, we regressed the network activity to the Bayesian state estimator *ŝ*_*t*_ and the belief threshold 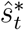 using the log-likelihood ratio, then calculated the decision variable as 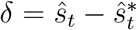.All regression models were trained on half of the trial data and tested on the remaining half.

### Bayesian Agent

The Bayesian agent was numerically simulated by performing a sequential update of the joint posterior distribution 𝒫_*t*_(*s, θ* | ***x***_≤*t*_) at each time step, following Eq. (4). The agent was assumed to be Bayes-optimal, possessing perfect knowledge of the world model, including the observation likelihood *p*(*x*_*t*_|*s*_*t*_, *θ*_*t*_) and the latent transition probabilities *P* (*s*_*t*_|*s*_*t*−1_) and *P* (*θ*_*t*_|*θ*_*t*−1_).

The agent initiated an action when the state estimate

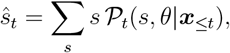

exceeded the optimal decision threshold,

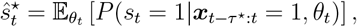

as defined in STAR Methods S.2.1. The optimal action time *τ*^⋆^ was determined by maximizing the expected reward,

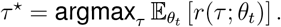

## A Supplemental information

**Figure S1:**
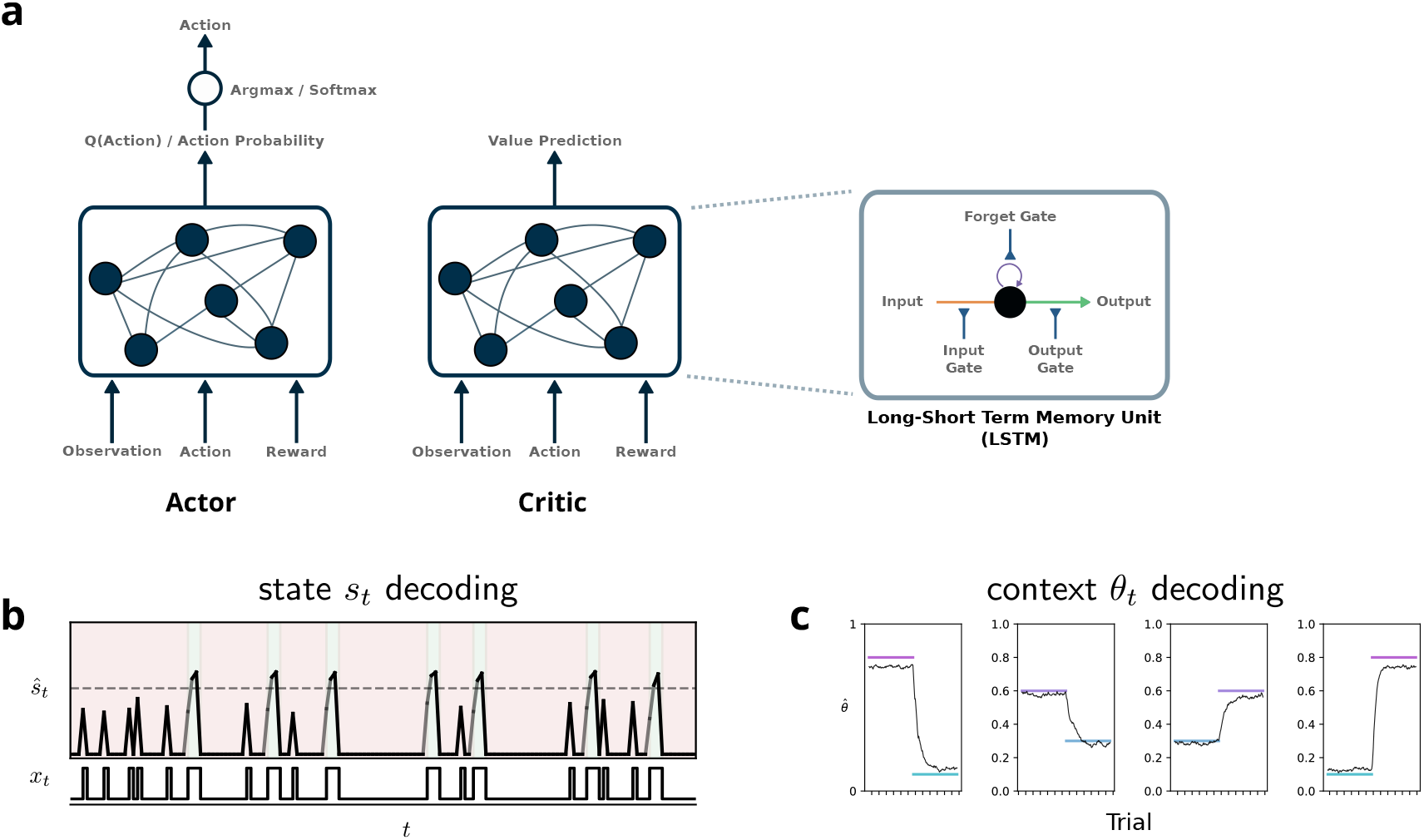
Network architecture and task-variable decoding. **(a)** Actor-critic and LSTM architecture. Actor and critic were each comprised of a Pytorch LSTM. **(b)** State decoding for example stimulus trajectory. Background color indicates ground truth *s*_*t*_. Dashed horizontal line indicates *ŝ*_*t*_ = 0.5 and line thickness corresponds to estimate accuracy. **(c)** Context decoding for example block transitions (averaged over several transitions). Colored lines indicate ground truth *θ*_*t*_.

### S.1 Principle component analysis

Here we show the PCA for the trained RNN and the untrained RNN which received an explicit context (Fig. S2a). This representation reveals that the representation is unsqueezed during training to give all contexts a similar dynamic range for the GO cues leading up to action. Stimulus statistics dominate the random dynamics, compressing the representation in high *θ*. An explicit context produces a context-dependent peak-action probability, however the representation is still entangled, preventing the readout from finding the optimal policy. The compressed representation in PC space corresponds to a saturation of the network DV (Fig. S2b) with only the trained network dedicating sufficient capacity to high *θ* for *Q*_*t*_ to reach values that support a deterministic policy.

### S.2 Details on analytical solution

#### S.2.1 Closed-form solution for reward rate in fixed context

Determining the optimal strategy for action requires estimating *P* (*s*_*t*_ | ***x***_≤*t*_) and thresholding on its value. Because *x*_*t*_ = 0 implies *s*_*t*_ = 0, it is sufficient to count cues from the last negative observation *x*_*t*−*τ*_ = 0, i.e. *P* (*s*_*t*_|***x***_≤*t*_) = *P* (*s*_*t*_|***x***_*t*−*τ*:*t*_ = 1, *x*_*t*−*τ*_ = 0), where we use the slicing notation *t*− *τ* : *t* to mean all times *t*′ fulfilling *t*− *τ < t*′≤ *t* (Matlab-style indexing). The waiting time *τ* hence forms the central component in the actor’s policy.

**Figure S2:**
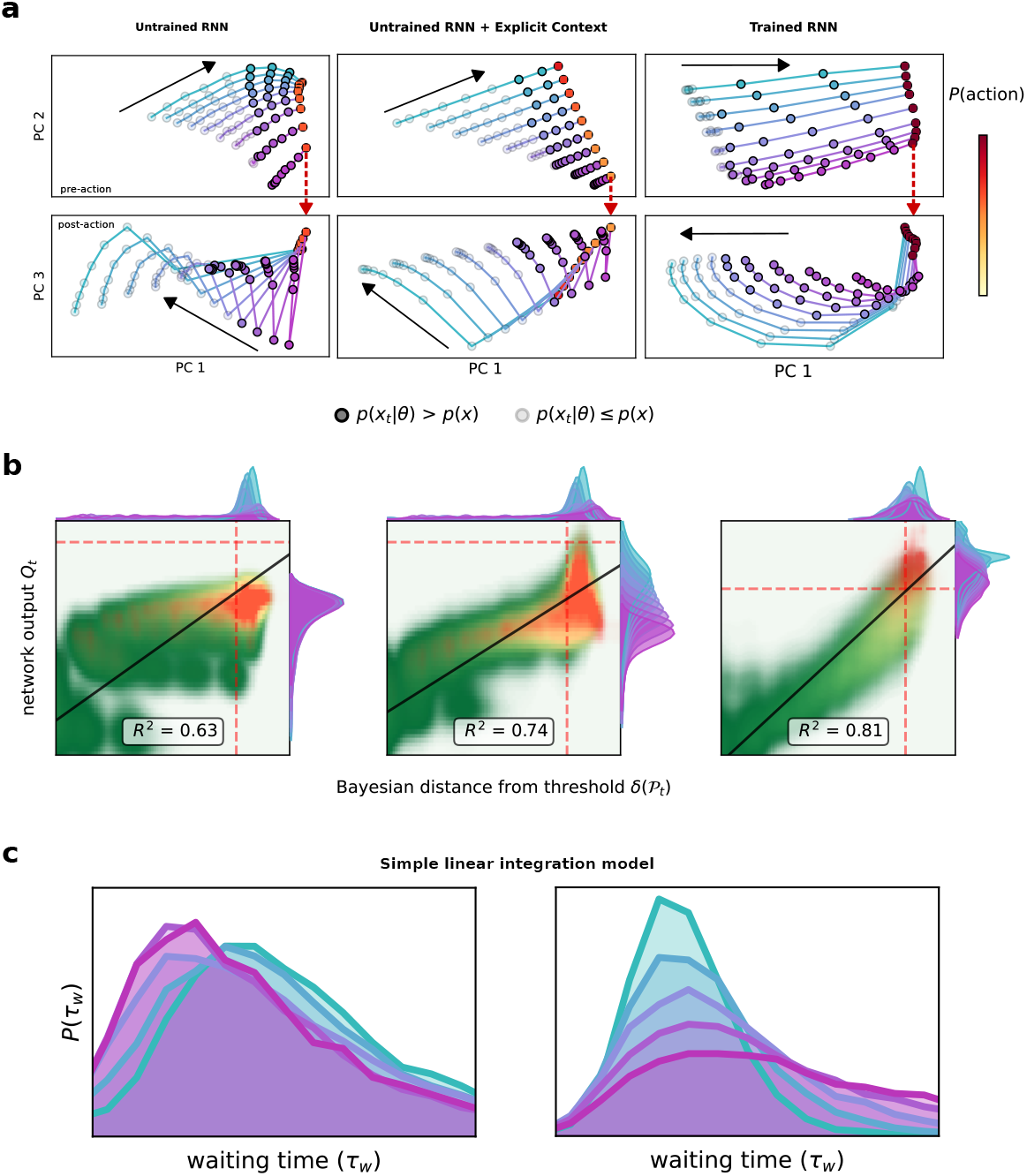
Network representations. **(a)** Projection of network activity on the first 3 PCs. Top rows: Pre-action trajectory in PC 1 and PC 2, for 9 steps leading up to action. Bottom rows-Post-action trajectory in PC 1 and PC 3, for 9 steps following action. Panels are arranged such that trial start to trial end follows a clockwise motion. Color of the action time circles indicates ⟨ *p*(action) ⟩_*t*|*θ*_ at action time. **(b)** Heatmap of *Q*_*t*_ vs *δ*_*t*_ during *x*_*t*_ = 1 for each network. *Q*_*t*_ of the readout from untrained RNNs saturates before crossing 0. *Q*_*t*_ of trained RNNs resembles *δ*(𝒫_*t*_) and crosses 0 for all *θ*. **(c)** Simple linear integration model replicates untrained network behaviors. Left-model behavior with *ϵ*_*θ*_ = 0 (*α* = 0.6, *β* = 6, *ϵ*_0_ = −7.3), right-model behavior with *ϵ*_*θ*_ = −3*θ* (*α* = 0.6, *β* = 10, *ϵ*_0_ = −9.4).

In the case where cues are faithful estimates of the latent state, 𝔼 [*x*_*t*_] = *s*_*t*_, their summed number will be increasing evidence that the latent state has switched. This idea has been formalized in sequential decision making, where the log-odds ratio 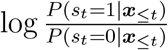 gets updated adaptively. In our setting, this translates to 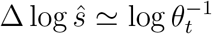 Herein, the inverse likelihood 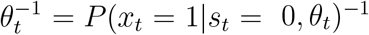 appears as the slope of the integrator, i.e., the reliability of the cues.

We further define auxiliary variables

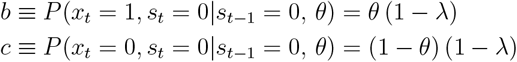

##### Transition after *τ* observations

In order to evaluate the reward rate in the main text, we need to calculate *P* (*s*_*t*_ = 1|***x***_≤*t*_, *θ*). Because an unsafe cue *x*_*t*_ = 0 always implies the unsafe state, it is sufficient to calculate *P*_*s*=1_ ≡*P* (*s*_*t*_ = 1| ***x***_*t*−*τ*:*t*_ = 1, *θ*), i.e. how likely the safe state is after *τ* suggestive observations, where we introduced *P*_*s*=1_ for brevity. To this end, we consider the complement *P*_*s*=0_ = 1 −*P*_*s*=1_. To evaluate this expression, we formulate a self-consistent relation,

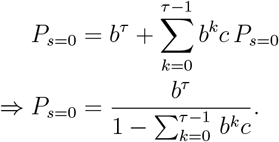

This equation considers the probability *b*^*τ*^ to have been unsafe the entire time. On top of this, the second sum considers all worlds where *k* observations *x* = 1 where followed by a faithful negative *x* = 0. By the Markov property of the world, these paths again reweight *P*_*s*=0_ itself. Resubstitution finally yields *P*_*s*=1_ = 1 − *P*_*s*=0_. To cross-check this relation, we simulated it via Monte Carlo sampling of the world.

##### Expected reward probability and action time

The average return when waiting for *τ* steps before acting is given by the probability the agent is acting in a safe state, *R* = *P* (*s*_*t*_ = 1 | *θ*, ***x***_*t*−*τ*:*t*_ = 1). The probability of being safe can be written explicitly as

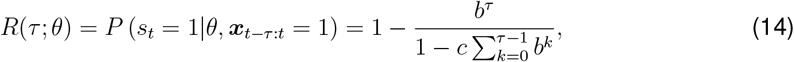

where *b*≡ (1 −*λ*)*θ* is the probability of a misleading *go* (i.e., the probability the state did not change times the probability of it being flipped), and *c* ≡ (1− *λ*)(1− *θ*) is the probability of observing a *nogo*.

To arrive at the final expression for the reward rate, the average time to reach *τ* cues is required, as well. We here extend the calculation presented by^76^ for the case of coin tosses with the latent context in our setting.

Let 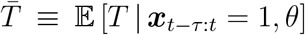 denote the expected action time, where *τ* indicates the period of consecutive cues *x*_*t*−*τ*:*t*_ = 1 preceding any action. Then, we arrive at a self-consistent relation

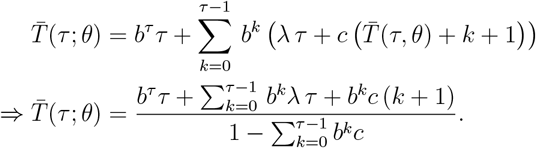

This considers all possible paths to get *τ* consecutive *x*_*t*−*τ*:*t*_ = 1: The first term is the probability of getting *τ unsafe* cues in *s*_*t*−*τ*:*t*_ = 0 right away. The sum then considers paths where *k* cues have been unsafe. From there on, a transition to *safe* happens with probability *a*, and all subsequent cues in this trial will be guaranteed to be faithfully *safe*. With probability *c* however, the world does not transition to *safe*. Again because of the Markovian structure, the world statistically indifferent from the initial time. Therefore, agent has to wait for 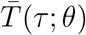 on expectation again, in addition to the cost *k* + 1 that it lasted to get to this point, with +1 accounting for the negative cue.

From this, the reward rate becomes

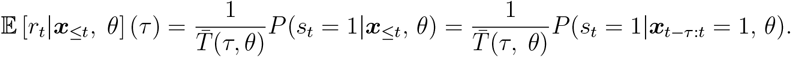

The maximizer *τ*^⋆^ of this expression defines the optimal policy in a context *θ*.

##### Latent variables require tracking joint density

In this section, we show that updating of the marginals of the joint belief cannot be done without holding track of the full joint distribution at the previous time step. This necessitates using the full joint update equation in the main text.

##### State

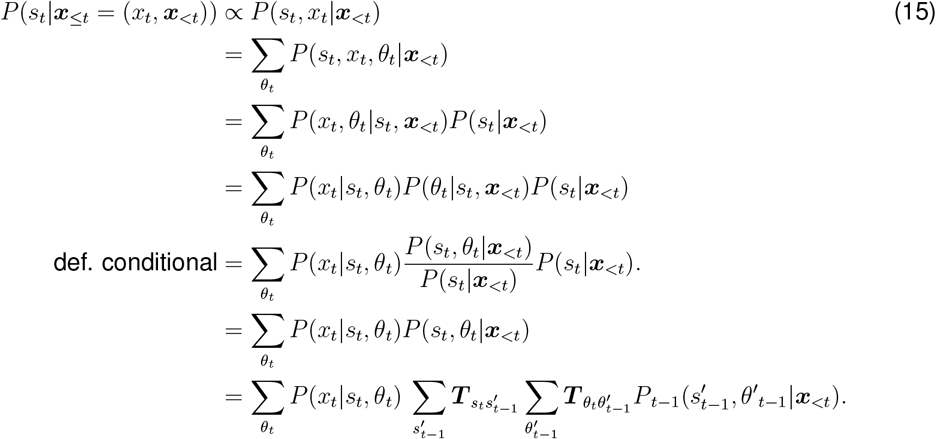

Importantly, in the third line, *P* (*s*_*t*_ |*θ*_*t*_, ***x***_*<t*_) ≠ *P* (*s*_*t*_ |***x***_*<t*_), in general. This can be seen from the fact that high noise will make the safe state more unlikely.

##### Context

This follows completely analogously

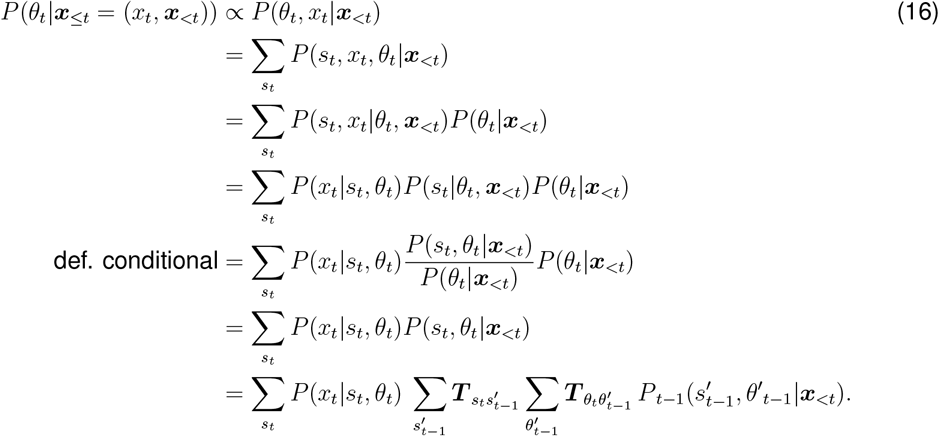

Note that these relations are completely equivalent to each other, up to a normalization.

In summary, this shows that the update of the marginals necessitates keeping track of the joint priors. Intuitively, this comes about in line 3, where the belief about the *joint* occurrence of *s*_*t*_, *θ*_*t*_ enters. Note that we can recover these relations by marginalization of the joint update in the main text, for example for the *s*_*t*_ update

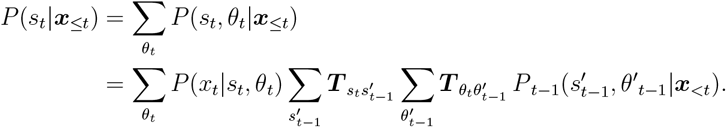

where we used the normalization of the transition matrix and the prior in the last two steps. This goes likewise for *P* (*θ*_*t*_ | ***x***_≤*t*_). The state and context estimators of the Bayesian agent were computed as averages with respect to the updated joint distribution from Eq. (4).

##### Bayesian inference without approximations

Because the transition functions were time-dependent, we made several approximations to make the inference process Markovian. The time-dependent transition functions without any approximations are as follows:

Time dependent ITI transition probabilities could be modeled as in^55^ with microstates (denoted as *s* = *k* for *k*∈ [−*u*_max_, …, −1]) where *u*_min_ and *u*_max_ are the minimum and maximum ITI durations. For each microstate the transition probability from ITI to unsafe is given by:

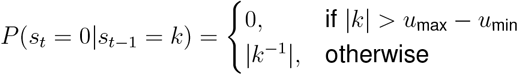

and the probability of transitioning to the next ITI microstate is simply given by

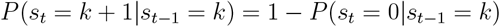

The fact that *θ* only changes at the end of an ITI is fully captured by:

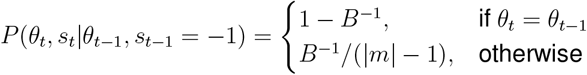

where *B* is the length of a block of constant *θ* and

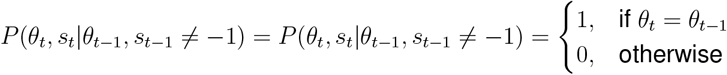

For the results in the main text we set *u*_min_ = *u*_max_ = 1 and approximated context transitions with stationary transition probabilities. This did not effect the performance of the Bayesian agent which remained near-optimal under both approximations.

